# Ewing Sarcoma Related protein 1 recognizes R-loops by binding DNA forks

**DOI:** 10.1101/2024.01.20.576463

**Authors:** Michelle A. Lay, Valery F. Thompson, Ajibola D. Adelakun, Jacob C. Schwartz

**Affiliations:** Department of Pharmacology, University of Arizona, Tucson, AZ 85724, USA; University of Arizona Cancer Center, Tucson, AZ 85724, USA; Department of Chemistry and Biochemistry, University of Arizona, Tucson, AZ 85724, USA; Department of Pharmaceutical Sciences, University of Arizona, Tucson, AZ 85724, USA

**Author notes:** Email:* M.A.L. V.F.T. A.D.A.

## Abstract

EWSR1 (Ewing Sarcoma Related protein 1) is an RNA binding protein that is ubiquitously expressed across cell lines and involved in multiple parts of RNA processing, such as transcription, splicing, and mRNA transport. EWSR1 has also been implicated in cellular mechanisms to control formation of R-loops, a three-stranded nucleic acid structure consisting of a DNA:RNA hybrid and a displaced single-stranded DNA strand. Unscheduled R-loops result in genomic and transcription stress. Loss of function of EWSR1 functions commonly found in Ewing Sarcoma correlates with high abundance of R-loops. In this study, we investigated the mechanism for EWSR1 to recognize an R-loop structure specifically. Using electrophoretic mobility shift assays (EMSA), we detected the high affinity binding of EWSR1 to substrates representing components found in R-loops. EWSR1 specificity could be isolated to the DNA fork region, which transitions between double- and single-stranded DNA. Our data suggests that the Zinc-finger domain (ZnF) with flanking arginine and glycine rich (RGG) domains provide high affinity binding, while the RNA recognition motif (RRM) with its RGG domains offer improved specificity. This model offers a rational for EWSR1 specificity to encompass a wide range in contexts due to the DNA forks always found with R-loops.

## INTRODUCTION

An R-loop is a structure formed by an RNA:DNA duplex inserted in stretch of double-stranded DNA (dsDNA), leaving the complementary strand to form a single-stranded DNA (ssDNA) loop ^1, 2^. Co-transcriptional R-loop formation occurs within highly transcribed genes when the nascent RNA leaving the polymerase binds its complimentary template DNA displacing the non-template strand ^2, 3^. Non-transcriptional R-loops can also be found in regions with guanine and cytosine (GC) rich sequences within the centromere and telomere ^1^. The interaction between an RNA:DNA hybrid is more stable compared to dsDNA. Formation of R-loops must be highly regulated to prevent transcription stalling and replication fork collision ^4, 5^. Failure to remove unscheduled R-loop may result in DNA damage and overall genomic instability ^6–8^.

Ewing Sarcoma is an aggressive pediatric bone cancer affecting 1 in 300,000 children in the United States, annually ^9^. These tumors are driven by a chromosomal translocation event resulting in a fusion protein, most commonly a fusion of the N-terminal low complexity (LC) domain of EWSR1 and C-terminal DNA binding domain of FLI1 protein, EWS-FLI1 ^10^. EWSR1 belongs to the FET (FUS, EWSR1, and TAF15) family of RNA binding proteins that are ubiquitously expressed and involved in many parts of RNA processing ^11, 12^. Other fusion proteins found in Ewing sarcoma can involve another C-terminal DNA-binding domain of the same ETS-family of transcription factors as FLI1 ^13, 14^. In rare instances of Ewing sarcoma, the LC domain of FUS might be substituted for that of EWSR1.

Recent studies have found EWSR1 can bind R-loops and has a role in their resolution, though much of the mechanism remains unknown ^3, 15–17^. In Ewing Sarcoma, the oncogenic fusion protein EWS-FLI1 has a dominant negative effect on the wildtype EWSR1 protein activity ^10^. This loss of EWSR1 activity leads to R-loops in Ewing Sarcoma cells being 4 times higher than any non-Ewing Sarcoma cell ^3^. In fact, these R-loops provide a transcriptional stress essential for Ewing sarcoma tumorigenesis ^3, 5, 18^. We have also recently reported a mechanism by which EWSR1-related protein, FUS, also mitigates R-loop accumulation ^19^.

We sought to investigate the mechanism of EWSR1 recognition of R-loop structures. By electrophoretic mobility shift assays (EMSA), we determined the binding affinity of EWSR1 to structures relating to R-loops. Subsequently, we investigated the contribution of domains in EWSR1 to its affinity for DNA. From this approach, we provide a model for EWSR1 to bind specifically to R-loop structures.

## MATERIALS AND METHODS

### Plasmid cloning

Wildtype (6x)His-MBP-EWSR1 gene was commercially synthesized in a pUC19 plasmid (Genscript). His-MBP-EWSR1 was inserted into a modified pGEX-6P-2 expression vector by restriction cloning with BsiWI and EcoRI restriction enzymes. Truncated EWSR1 constructs were created by PCR-based cloning from the His-MBP-EWSR1 expression vector using inverse PCR with Phusion DNA polymerase (New England Biolabs, F530S) then DpnI digest (New England Biolabs, 101228-926). The linear PCR product was then re-circularized with In-Fusion Cloning Master Mix (Takara Bio, 638948) according to manufacturer’s protocol.

### Recombinant protein expression and purification

Expression vectors for full-length and truncated His-MBP-EWSR1 protein were produced essentially as previously published ^20^. Transformed BL21(DE3) *E. Coli* (NEB, C2527I) were grown on LB (Luria-Bertani) agar plate (0.1 mg/mL ampicillin) and a colony selected was then grown overnight in 10 mL of LB media at 37° C (0.1 mg/mL ampicillin) with shaking. After inoculating 1 L of LB media (0.1 mg/mL ampicillin), the cultures were grown until OD_600_ 0.6-0.8. Protein expression was induced with 1 mM IPTG and expression continued overnight at 17° C. Cells were collected by centrifugation (4000×g) at 4° C for 20 minutes then resuspended in EWS buffer (1M Urea, 1M KCl, 50 mM Tris-HCl pH = 8.0, 10 mM Imidazole, 5% glycerol) supplemented by 1% NP-40 and 1.5 mM β-mercaptoethanol. Lysates were prepared by sonication, cleared by spinning at 20,000×g for 20 minutes and 4° C, then incubated 1 hour with Ni-NTA Sepharose beads (Cytiva, 17-5318-01) at 4° C. Ni-NTA beads were washed 4x in EWS buffer. Protein was eluted from Ni-NTA beads using elution buffer (1M Urea, 1M KCl, 50 mM Tris-HCl pH = 7.4, 250 mM Imidazole) for 20 minutes at room temperature. Protein purity was assessed by SDS-PAGE and protein stored at room temperature up to 1 month.

| Gene | Backbone | Tag | Amino Acids |
| --- | --- | --- | --- |
| H-MBP-EWSR1 | pGEX-6P-2 | N-H-MBP | M1-Y656 |
| del-LC EWSR1 | pGEX-6P-2 | N-H-MBP-MASTD | G290-Y656 |
| R<sub>1</sub>-RRM-R<sub>2</sub> EWSR1 | pGEX-6P-2 | N-H-MBP-MASTD | G290-V515 |
| R<sub>2</sub>-ZnF-R<sub>3</sub> EWSR1 | pGEX-6P-2 | N-HMBP-MASTD | M454-Y656 |

### Radiolabeled Nucleic Acid Substrate Preparation

Single stranded nucleic acid strand was chemically synthesized from SigmaAldrich. DNA substrates were 5’ terminally labelled with T4 Polynucleotide Kinase (NEB, M0201S) and ATP-γ-^32^P (PerkinElmer, NEG002A100UC) following manufacturer’s protocol. Unincorporated free ATP was removed from reaction product using G-25 column (Cytiva, 27-5325-01). Equimolar concentration of oligos in annealing buffer (60 mM KCl, 1x PBS, and 0.2 mM MgCl_2_) were annealed by incubation at 95°C (5 min) then reduced to 25° C by step-wise intervals of 10° C (6 min/step). The dsDNA-88 substrate was produced by PCR amplification of R-loop-88-Fwd oligo with DNA Phusion polymerase.

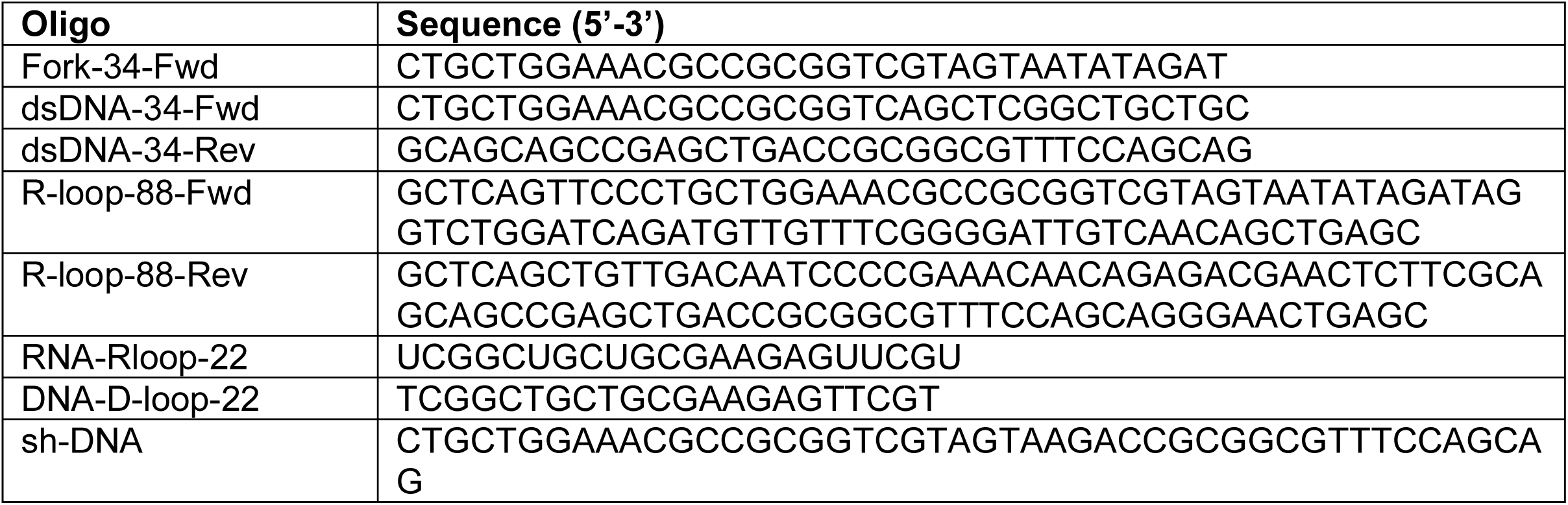

### Electrophoretic mobility shift assay (EMSA)

Nucleic acid substrates labelled by γ-^32^P (20000 cpm) were incubated with protein in 20 μL reaction buffer (40 mM Tris-HCl pH = 8.0, 5 mM DTT, 1.5 mM MgCl_2_) for 30 minutes at room temperature. Binding was assessed using electrophoresis (100V for 50 min) with a 7.5% acrylamide Tris-Borate-EDTA (TBE) native gel at room temperature, which was then transferred to Nylon+ membrane (Cytiva, 45-000-850) at 50 V for 1 hour (4° C). Radiodensitometry analysis was made by overnight exposure to phosphor screen (Cytiva, 28-9564-75) and imaged by Typhoon-FLA 9000, then quantified with ImageLab. Curve fits and binding constants were determined using MyCurveFit software.

## RESULTS AND DISCUSSION

### EWSR1 preferentially binds DNA structure containing loops

To assess EWSR1 specificity to bind R-loop structures, we designed and hybridized 88-nt long DNA oligomers with complementary sequences surrounding an internal 28 nt long non-complementary sequence. RNA was annealed to one strand of the loop region to form the R-loop (R-loop-88). We performed EMSA analysis to measure dissociation constants (K_D_) of binding for recombinant expressed and purified EWSR1 fused by an N-terminal 6xHis and a maltose-binding protein tag for solubility, H-MBP-EWSR1.

With increasing protein concentration, EMSA did show H-MBP-EWSR1 form a complex with R-loop-88 (**Figure 1A**). A strong binding affinity was evidenced by an equilibrium dissociation constant, K_D_, in the low nanomolar range, 72 ± 28 nM (**Figure 1B, Supplemental Figure 1A**). Multiple species of bound complexes were observed, suggesting a multimer of EWSR1 bound to the R-loop. To test the importance of the RNA:DNA hybrid, the RNA strand bound to the loop was replaced by a DNA oligonucleotide (D-loop-88). H-MBP-EWSR1 bound D-loop-88 with a similar K_D_, 55 ± 8.3 nM, to that of the R-loop (**Figure 1A–B, Supplemental Figure 1B**). EWSR1 also bound an empty loop (Empty-Loop) with a similar affinity, K_D_ = 46 ± 15 nM (**Figure 1A–B, Supplemental Figure 1C**). These data showed that high affinity binding of EWSR1 depended neither on the identity nor presence of a third nucleic acid strand in DNA loop.

**Figure 1:**
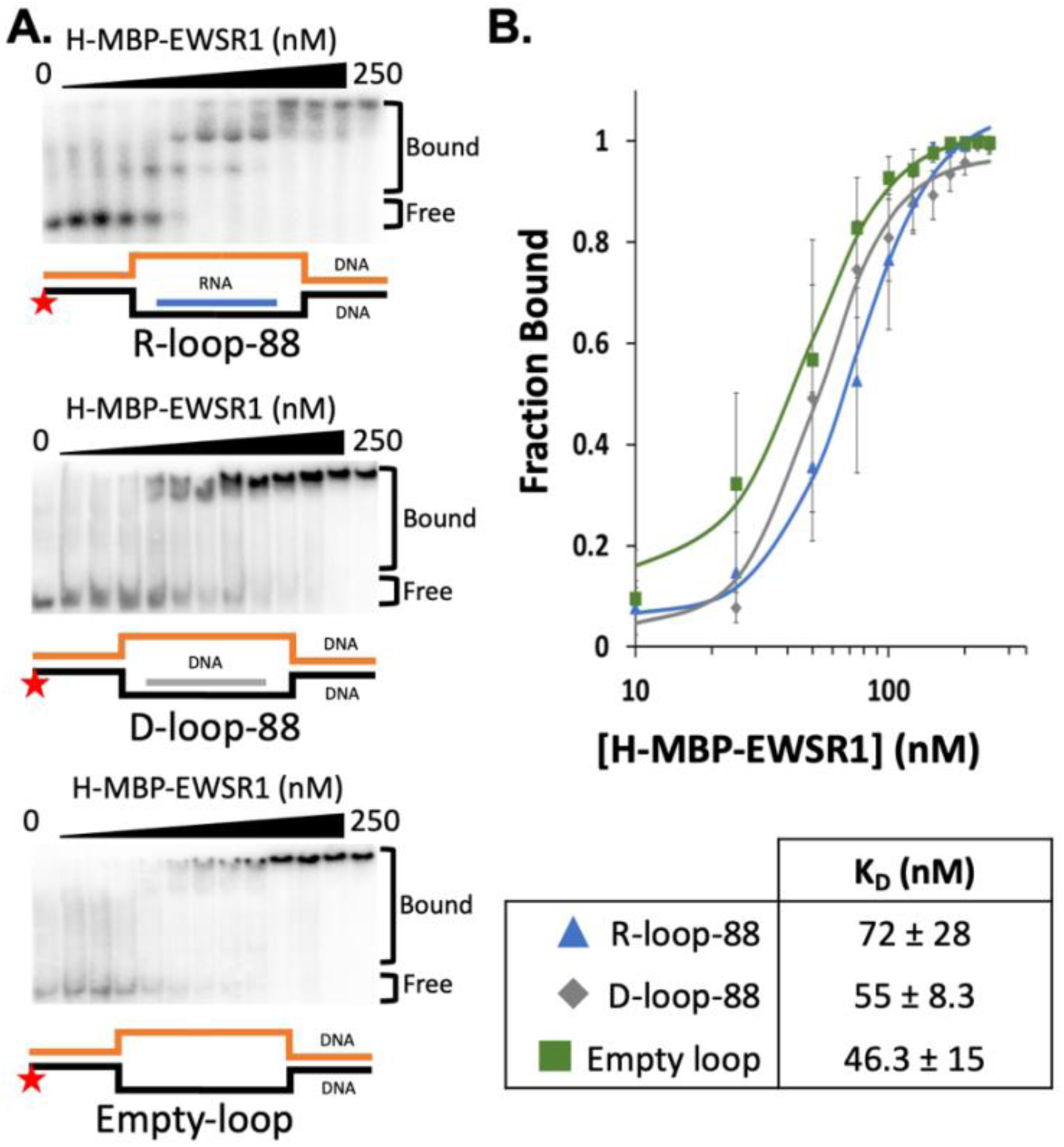
H-MBP-EWSR1 preferentially binds DNA loop structures. (**A**) Trace amount of radiolabeled nucleic acid substrates are incubated with increasing concentration of H-MBP-EWSR1. Binding was observed through electrophoretic mobility shift assay (EMSA). (**B**) Fraction of nucleic acid bound over total nucleic acid substrate was plotted against H-MBP-EWSR1 concentration. Amount of H-MBP-EWSR1 needed to shift half of the substrate is measured as the binding affinity (K_d_) of H-MBP-EWSR1 and that substrate.

To compare EWSR1 binding specificity for loop structures, we synthesized three new structures for study: an 88 bp double stranded DNA (dsDNA-88), single-stranded DNA (ssDNA-88), or RNA:DNA hybrid alone (Hybrid-88). EWSR1 showed strong binding affinity for ssDNA-88, K_D_ = 62 ± 13 nM (**Supplemental Figure 1D-E**). We measured 2x weaker affinity for EWSR1 binding to dsDNA-88, K_D_ = 103 ± 5 nM, relative to loop structures (**Supplemental Figure 1D, 1F**). EWSR1 showed the least affinity for Hybrid-88 with a K_D_ of 137 ± 2 nM (**Supplemental Figure 1D, 1G**).

### Both double and single stranded DNA region is needed for EWSR1 tight binding

While EWSR1 showed highest affinity for both loop and single-stranded DNA, we wanted a more unambiguous rationale for binding specificity. Because of its length, ssDNA-88 could fold into a variety of structures resembling other portions of the loops. One structure common to the three loop structures is a DNA fork ^1^. We prepared a 34 nucleotide DNA fork (D-fork-34) with a 20 bp region and 14 nt single-stranded region of non-complementary sequence (**Figure 2A**, **Supplemental Figure 2A**). EWSR1 fully shifted the D-fork-34 with K_D_ = 59 ± 5 nM (**Figure 2A–B**, see also **Table 1**). EWSR1 bound a 34 bp dsDNA (dsDNA-34) with an intermediate affinity, K_D_ = 126 ± 15 nM (**Figure 2A–B, Supplemental Figure 2B**). EWSR1 showed relatively weak affinity for a 34-nucleotide ssDNA (ssDNA-34), K_D_ = 216 ± 9 nM (**Figure 2A–B, Supplemental Figure 2C**).

**Figure 2:**
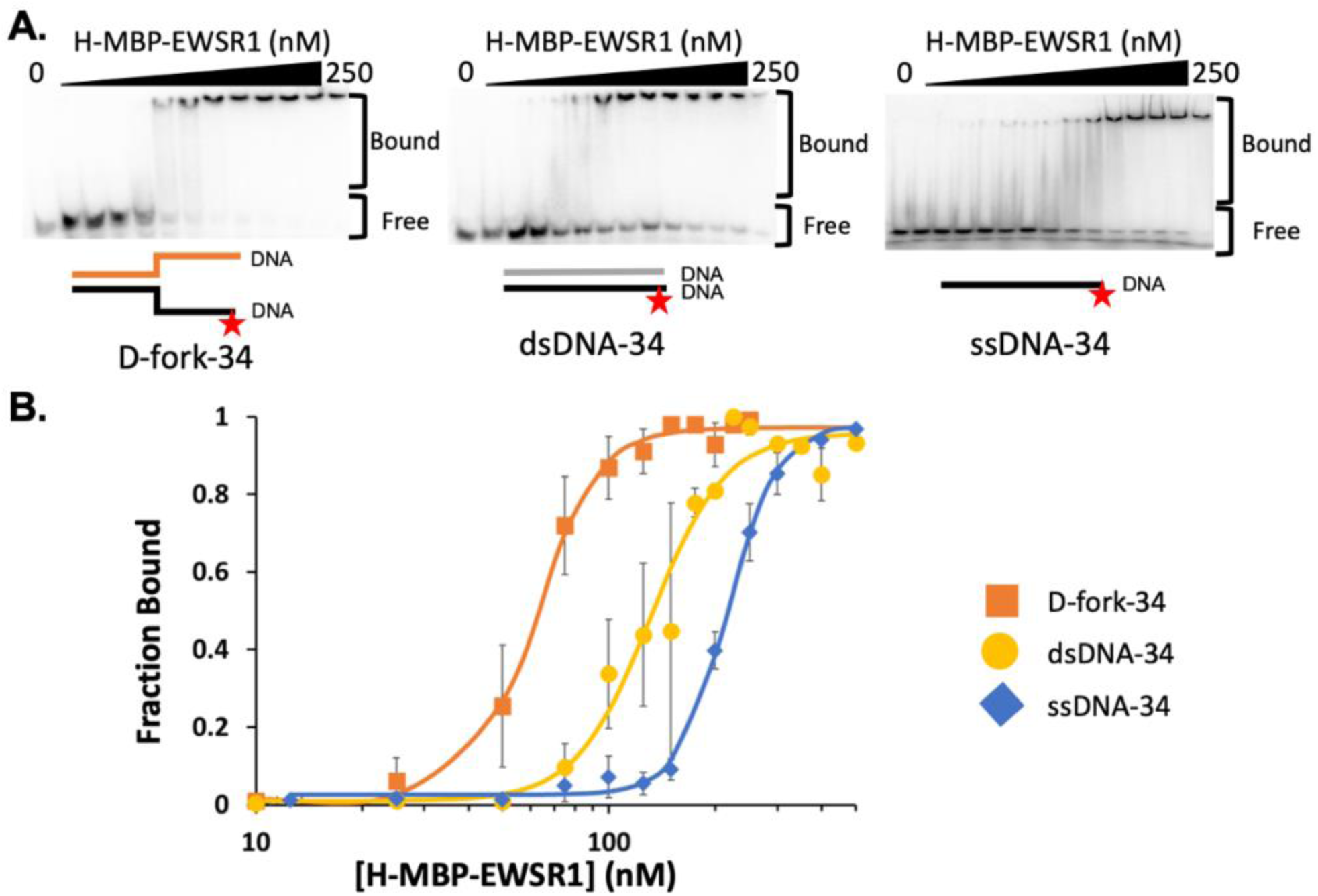
H-MBP-EWSR1 binds structures containing junctions. (**A**) Binding ability of H-MBP-EWSR1 with 34 nucleotides DNA constructs Fork-34, dsDNA-34, and ssDNA-34 observed through EMSA. (**B**) Band shift from EMSA was quantified to measure binding affinity as K_d_

**Table 1:**
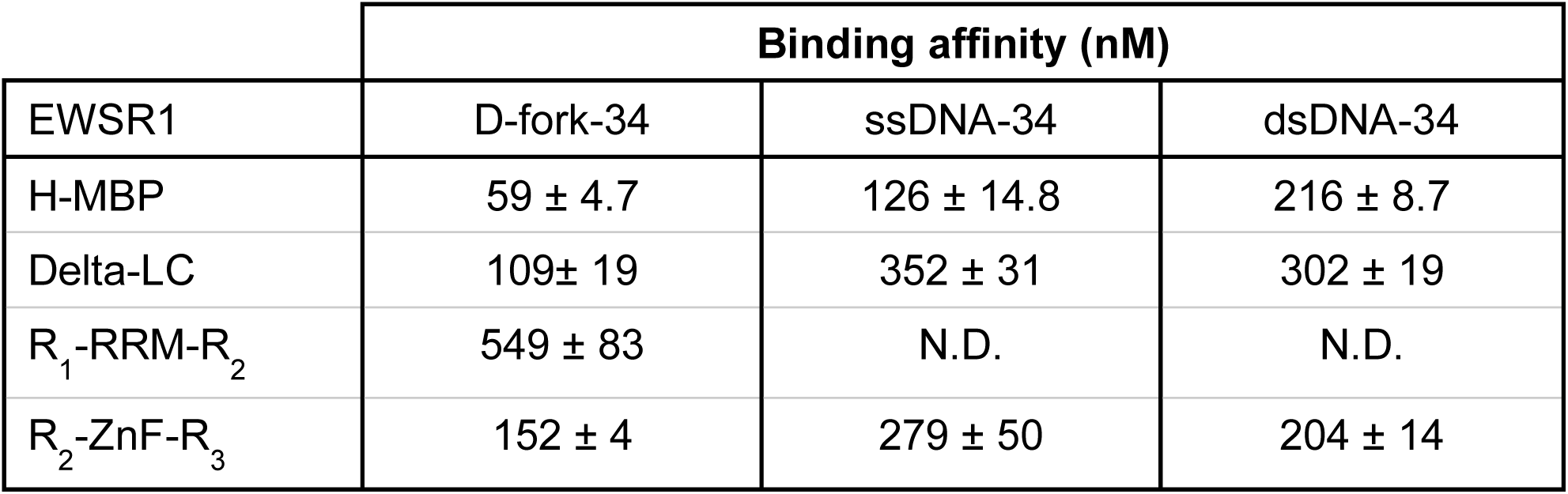
A summary of the binding affinity between EWSR1 constructs (wildtype, delta LC, R_1_-RRM-R_2_, and R_2_-ZnF-R_3_) with 34 nt DNA constructs (Fork-34, ssDNA-34, and dsDNA-34)

Lastly, we considered a DNA hairpin structure, which partially resembles a fork but the flexibility of the single stranded region is more constrained. We prepared a 47 nt stem loop structure that when folded contains a 20 bp region and a 7 nt loop (sh-DNA). By EMSA, we found the K_D_ of binding for EWSR1 to sh-DNA stem loop to be 236 ± 5 nM, making the hairpin DNA the least preferred structure (**Supplemental Figure 2D)**.

### Domain contributions of EWSR1 to DNA binding

The LC domains of FET proteins are important for their homo-typic interactions that form higher order assemblies ^10, 21, 22^. We asked whether LC domain of EWSR1 contributes to binding specifically and tightly to DNA. We deleted residues D6 to G289 from EWSR1 (del-LC-EWSR1, **Figure 3A**) and found del-LC-EWSR1 bound D-fork-34 with moderate affinity and a K_D_ of 109 ± 19 nM, almost twice that of H-MBP-EWSR1 (**Figure 3B–C, Supplemental Figure 3A)**. Binding affinity of del-LC-EWSR1 for ssDNA and dsDNA, K_D_ of 302 ± 19 nM and 352 ± 31 nM, respectively (**Figure 3B–C, Supplemental Figure 3A**). Together these results indicate LC domain does have a role in supporting high affinity binding to DNA (**Table 1**).

**Figure 3:**
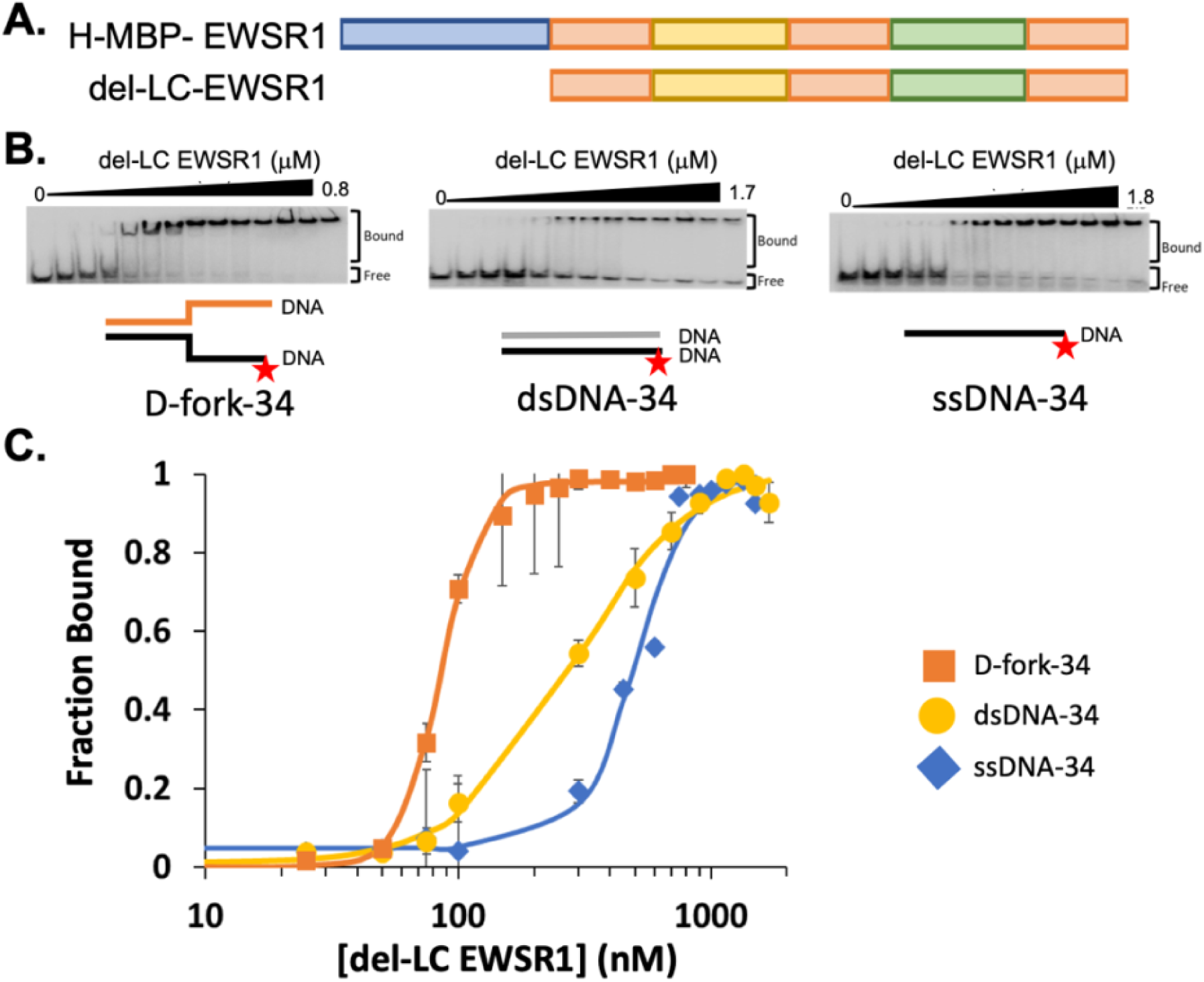
Contribution of LC domain of EWSR1 to nucleic acid binding ability. (**A**) Domain structures of del-LC EWSR1 truncation. (**B**) EMSA of del-LC with ssDNA-34 (left), dsDNA-34 (middle), and Fork-34 (right). (C). Band shift from EMSA was quantified to measure binding affinity as K_d_

We next explored the contribution of the nucleic acid binding domains in EWSR1 to DNA binding. We created two truncations of EWSR1: R_1_-RRM-R_2_ (a.a. 290-515) and R_2_-ZnF-R_3_ (a.a. 454-656) (**Figure 4A**). Very weak binding was found for R_1_-RRM-R_2_ to the D-Fork-34 structure (K_D_ = 549 ± 83 nM, **Figure 4B–C, Supplemental Figure 4A**). The binding affinity of R_1_-RRM-R_2_ to ssDNA-34 and dsDNA-34 could not be determined (**Figure 4B–C, Supplemental Figure 4B–C**). R_2_-ZnF-R_3_ showed a higher affinity for DNA substrates than R_1_-RRM-R_2_. R_2_-ZnF-R_3_ bound tightest to D-fork-34, KD = 152 ± 4 nM (**Figure 4B, 4D, Supplemental Figure 4D**). R2-ZnF-R3 bound with only slightly less affinity to ssDNA-34 (204 ± 14 nM) and dsDNA-34 (279 ± 50 nM) relative to D-fork-34 (**Figure 4B, 4D, Supplemental Figure 4E–F**). In comparison, the affinity of R_1_-RRM-R_2_ for DNA was weak but R_2_-ZnF-R_3_ showed less than a 2-fold difference in K_D_ for D-fork-34 and ssDNA or dsDNA, indicating less specificity than H-MBP-EWSR1 or del-LC-EWSR1 (**Table 1**).

**Figure 4:**
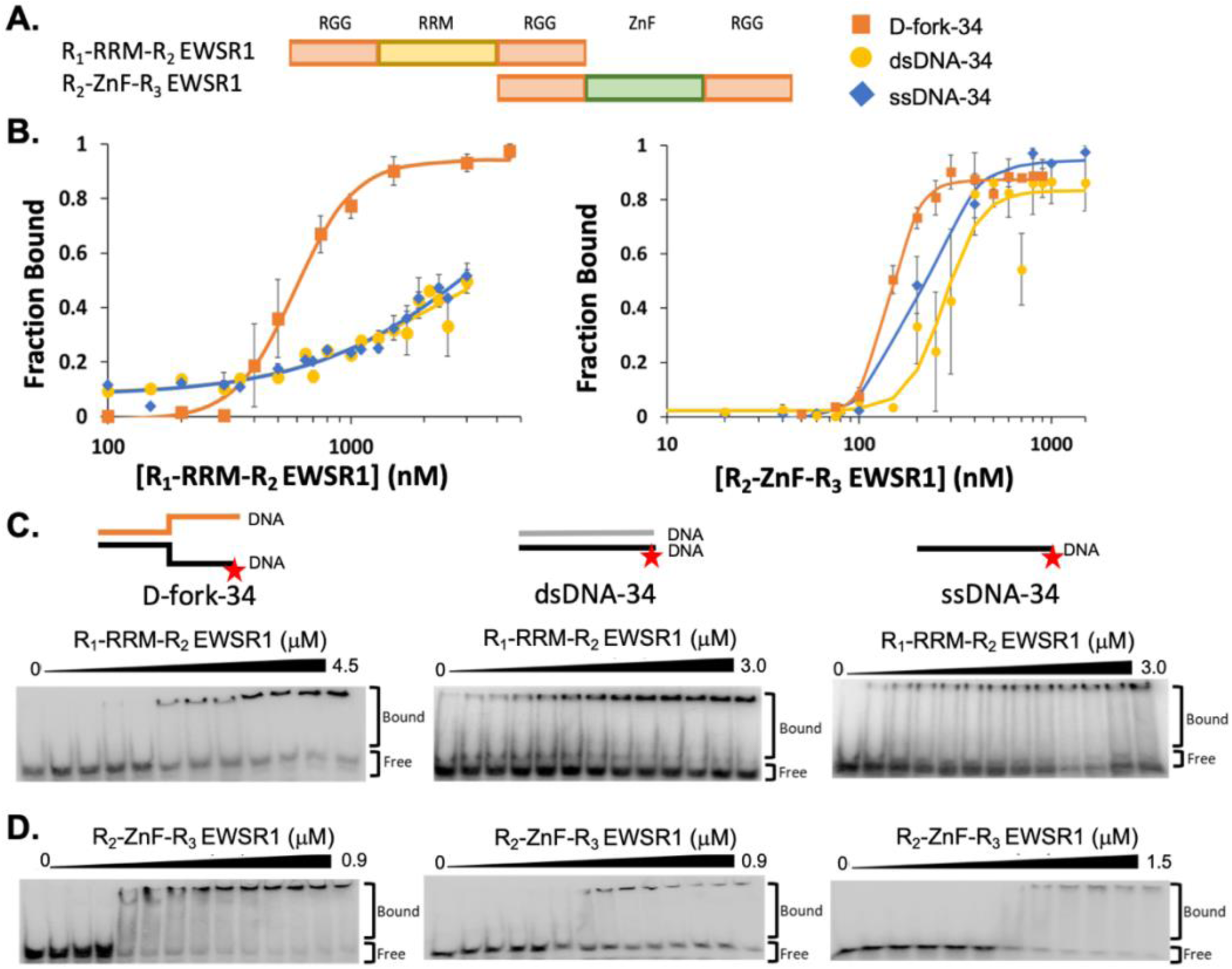
EWSR1 truncations to study the contribution of EWSR1 RNA binding domains to DNA binding. (**A**) Domain structures R_1_-RRM-R_2_ and R_2_-ZnF-R_3_ EWSR1. (**B**) EMSA quantified to measure the binding affinity of EWSR1 truncations with DNA substrates. (**C,D**) EMSA performed with R_1_-RGG-R_2_ and R_2_-RRM-R_3_ EWSR1 mutants with ssDNA-34 (left), dsDNA-34 (middle), and Fork-34 (right).

## CONCLUSION

In this study, we investigated EWSR1 binding and specificity to R-loop structures. We found that EWSR1 can bind an R-loop with a high affinity, but is also able to bind a D-loop and empty loop with similar affinity. The identity or presence of a invading nucleic acid in an R-loop is not necessary for EWSR1 recognition. We show that a DNA fork, which is found at the boundary of a loop and contains dsDNA and ssDNA, is sufficient for tight binding by EWSR1 (< 100 nM affinity). Exploring contributions of domains in EWSR1 revealed that the ZnF and its flanking RGG domains contribute the most to high affinity DNA binding. Although the RRM and flanking RGG domains bind poorly, these domains are necessary for high specificity between DNA fork, dsDNA, or ssDNA. The LC domain also influences DNA binding by EWSR1. It is likely the LC domain provides oligomerization to enhance binding to DNA ^21–24^.

Our study finds similar high affinity binding of EWSR1 to DNA and R-loops as in previous reports ^16, 17^. These are also comparable to the highest affinities observed among FET protein and a nucleic acid target ^12, 25^. The typical RNA binding domains contain more than one RNA binding domain to increase RNA binding specificity and affinity ^26, 27^. The two most common RBDs are the RRMs and RGGs which can be found in up to 20% of RBPs ^28, 29^. The RRM of FUS recognizes the single-stranded portion of an RNA stem loop, but we found EWSR1 had low affinity for a DNA stem loop ^30^. While most RRM domains bind RNA, some exceptions of DNA binding and processing have been reported. Interaction between RRM of hnRNPA1 with ssDNA has been confirmed with a crystal structure ^31, 32^. The RGG domain in hnRNP A1 was found to destabilize DNA G-quadraplexes, the RRM binds and stabilizes the now resolved ssDNA ^33, 34^. HNRPDL (JKTBP), made up of 2 RRM domains and 1 RGG domain, can bind both single and double stranded DNA ^35^. Protein-DNA interaction in hnRNP A1 and HNRPDL are both important for transcription regulation ^33, 35^. Interaction between RGG and DNA G-quadraplexes has been observed in EWSR1 ^17, 36, 37^. This raises the possibility that when G-quadruplex structures are present in the single stranded DNA of the R-loop, EWSR1 recruitment can be improved ^1, 18^.

R-loops occur naturally in the nucleus and are typically transient ^4, 38^. Their repair involves recruitment of so many proteins, a reasonable prediction is that a molecular scaffold like that formed by the assembly of EWSR1 oligomers can be required for efficient activity ^22, 39^. However, unscheduled resolution of R-loops can result in head on collision with replication fork and transcription stalling which eventually leads to overall genomic stability found in many diseases ^2, 4, 5^. In Ewing sarcoma, wildtype EWSR1 is essentially inactive, and the number of R-loops is up to 4x higher compared to normal healthy cells ^3, 16, 17^. Our data provides a mechanism to involve EWSR1 in R-loop resolution through a general approach to recognize R-loops.

## ACKNOWLEDGEMENTS

This work was supported by the National Institutes of Health [CA259570 to J.C.S.]; the American Cancer Society [RSG-18-237-01-DMC to J.C.S.]; and a training fellowship from the University of Arizona Cancer Center to M.A.L. The content is solely the responsibility of the authors and does not necessarily represent the official views of the National Institutes of Health.

---

**Supplemental Figure 1:**
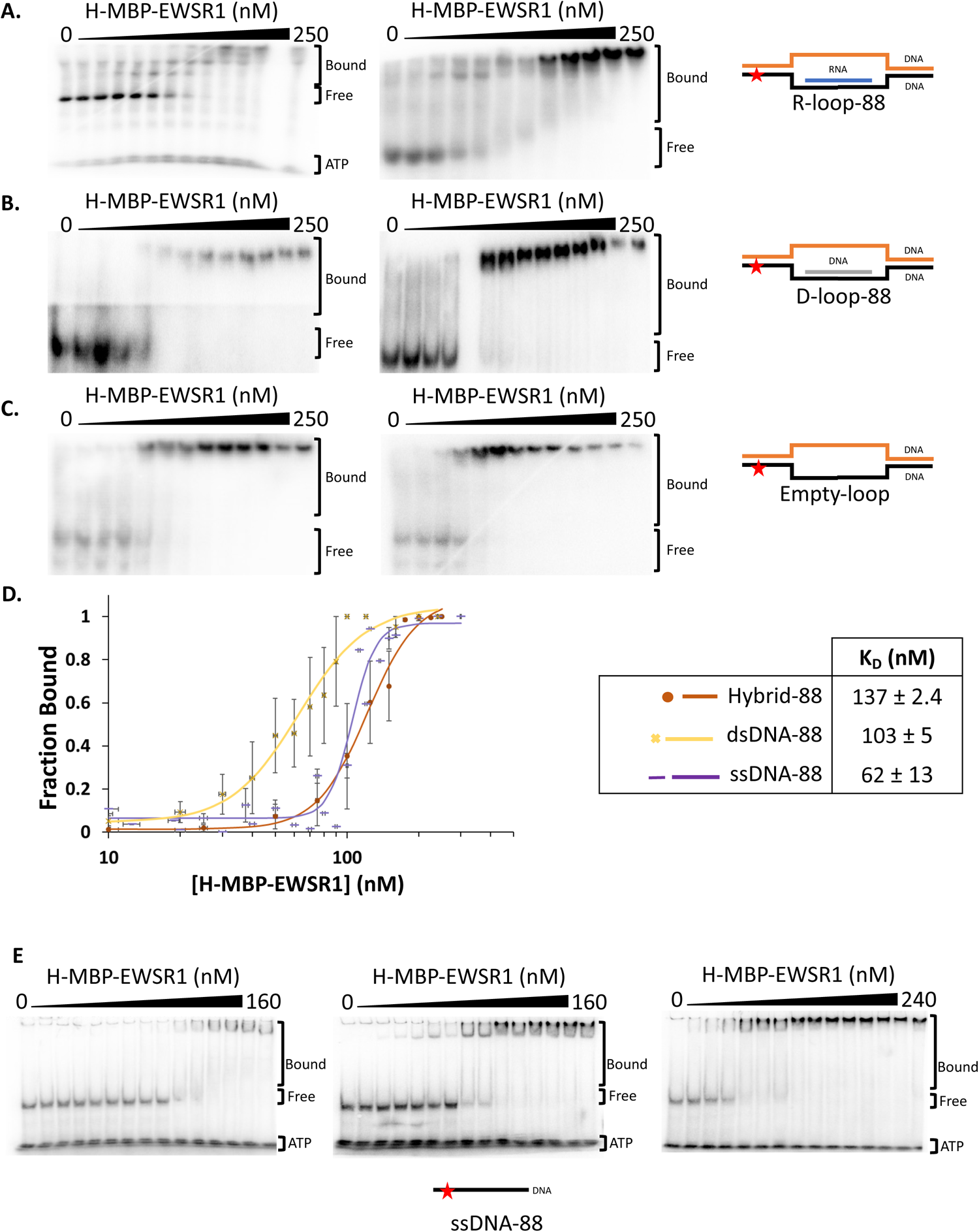
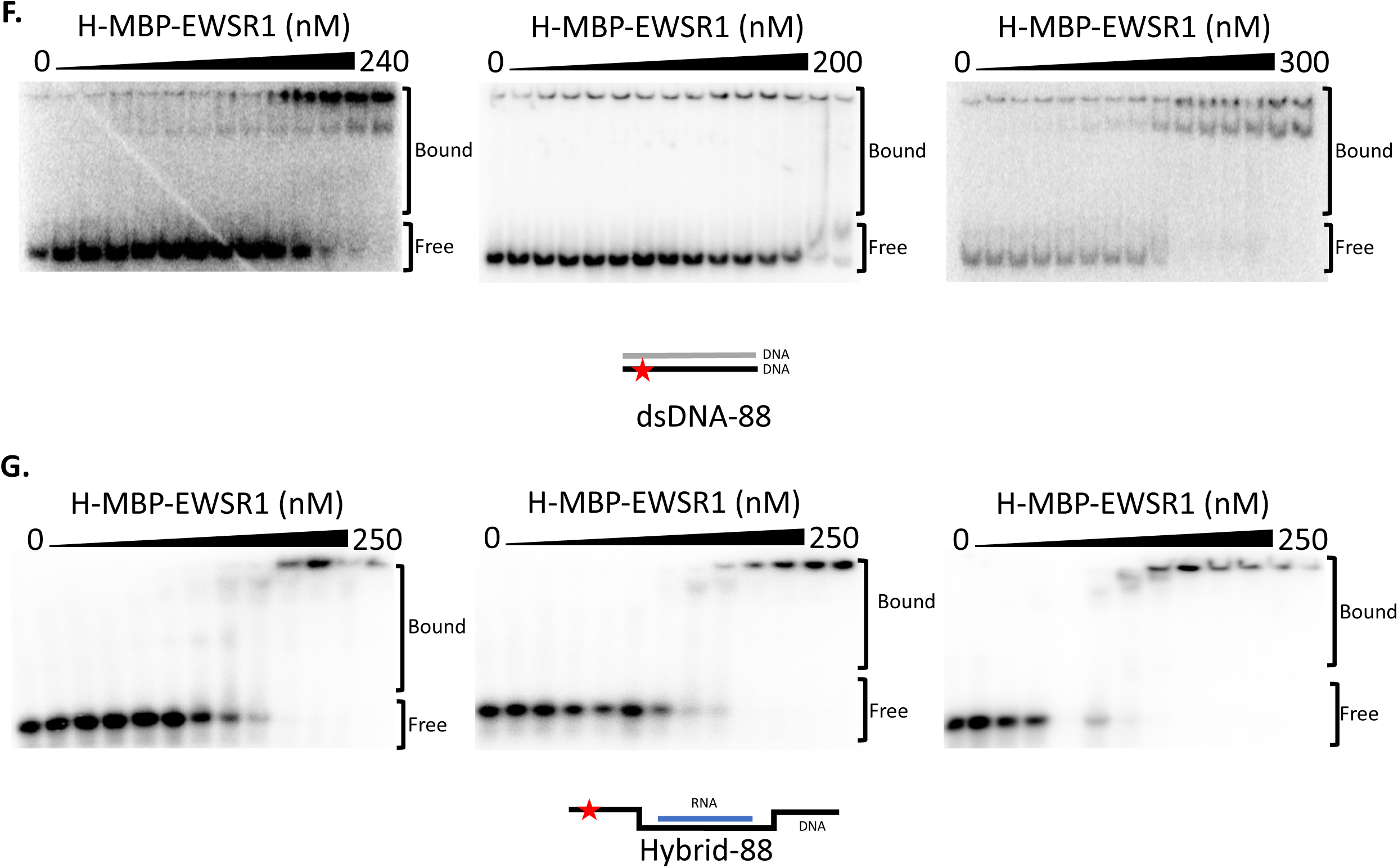
EMSA replicates for binding of H-MBP-EWSR1 to (A) R-loop-88 (B). D-loop-88 and (C) Empty loop. (D) Plot of average fraction of nucleic acid bound for RNA:DNA Hybrid-88 (E), dsDNA-88 (F), and ssDNA-88 (G). Error is shown as standard error about the mean, N = 3.

**Supplemental Figure 2:**
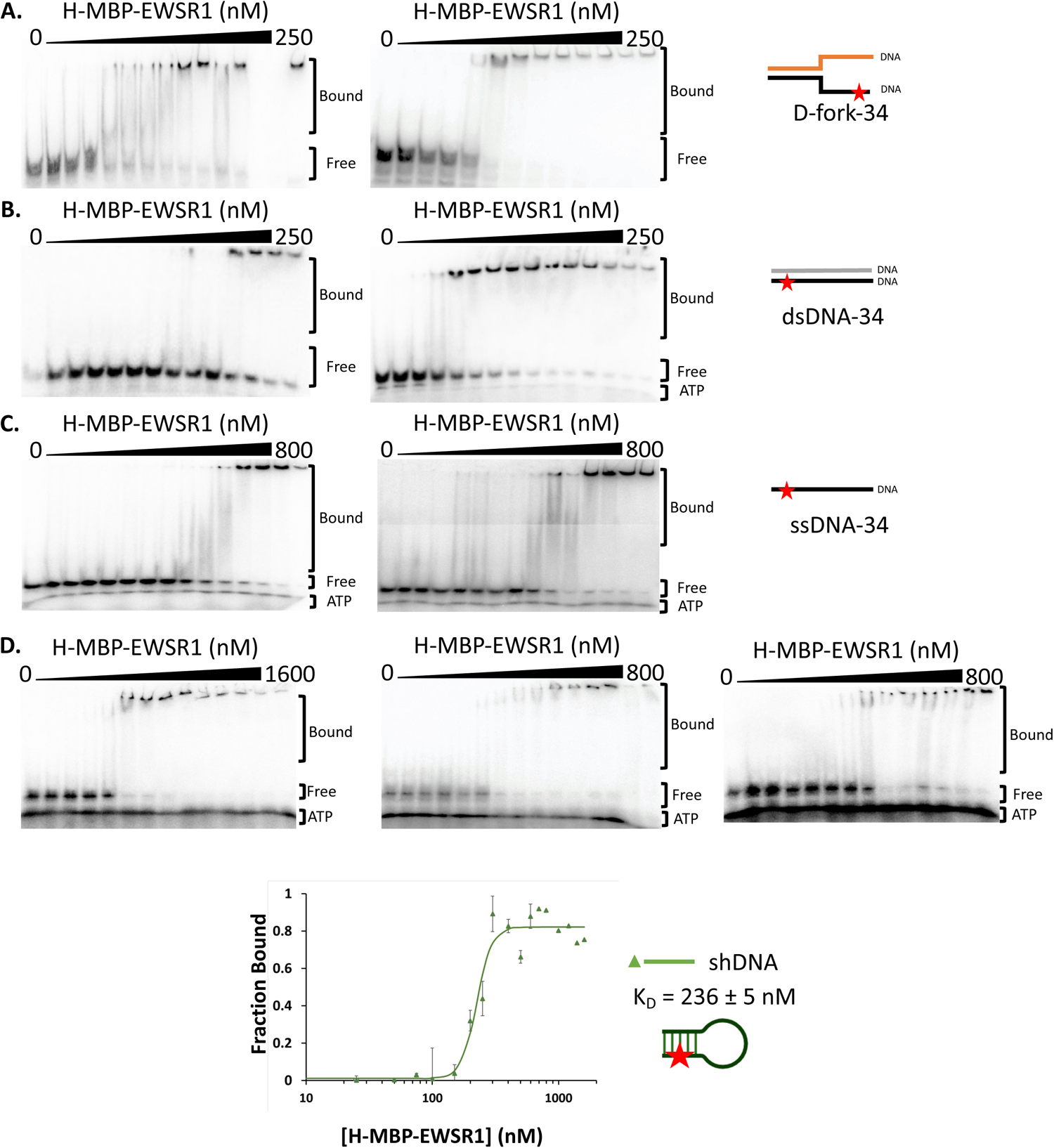
EMSA analysis of H-MBP-EWSR1 binding to (**A**) D-fork-34, (**B**) dsDNA-34, (**C**) ssDNA-34, and (**D**) shDNA. Average fraction of sh-DNA bound was plotted against H-MBP-EWSR1 concentration and the averaged equilibrium dissociation constant, K_D_, determined. Error is shown as standard error about the mean, N = 3.

**Supplemental Figure 3.**
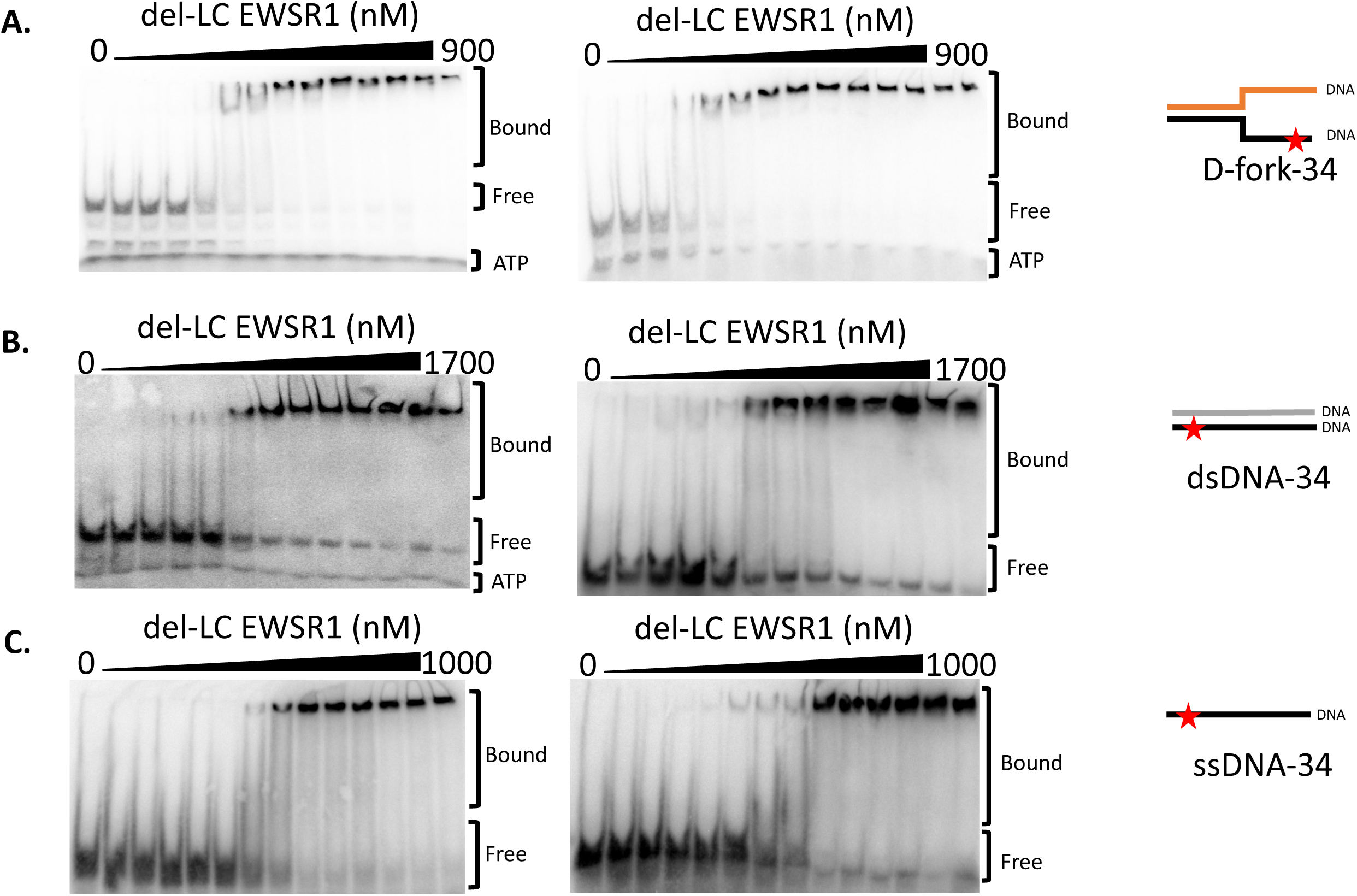
EMSA replicates to show binding of del-LC-EWSR1 and (A) D-fork-34, (B) dsDNA-34, and (C) ssDNA-34.

**Supplemental Figure 4.**
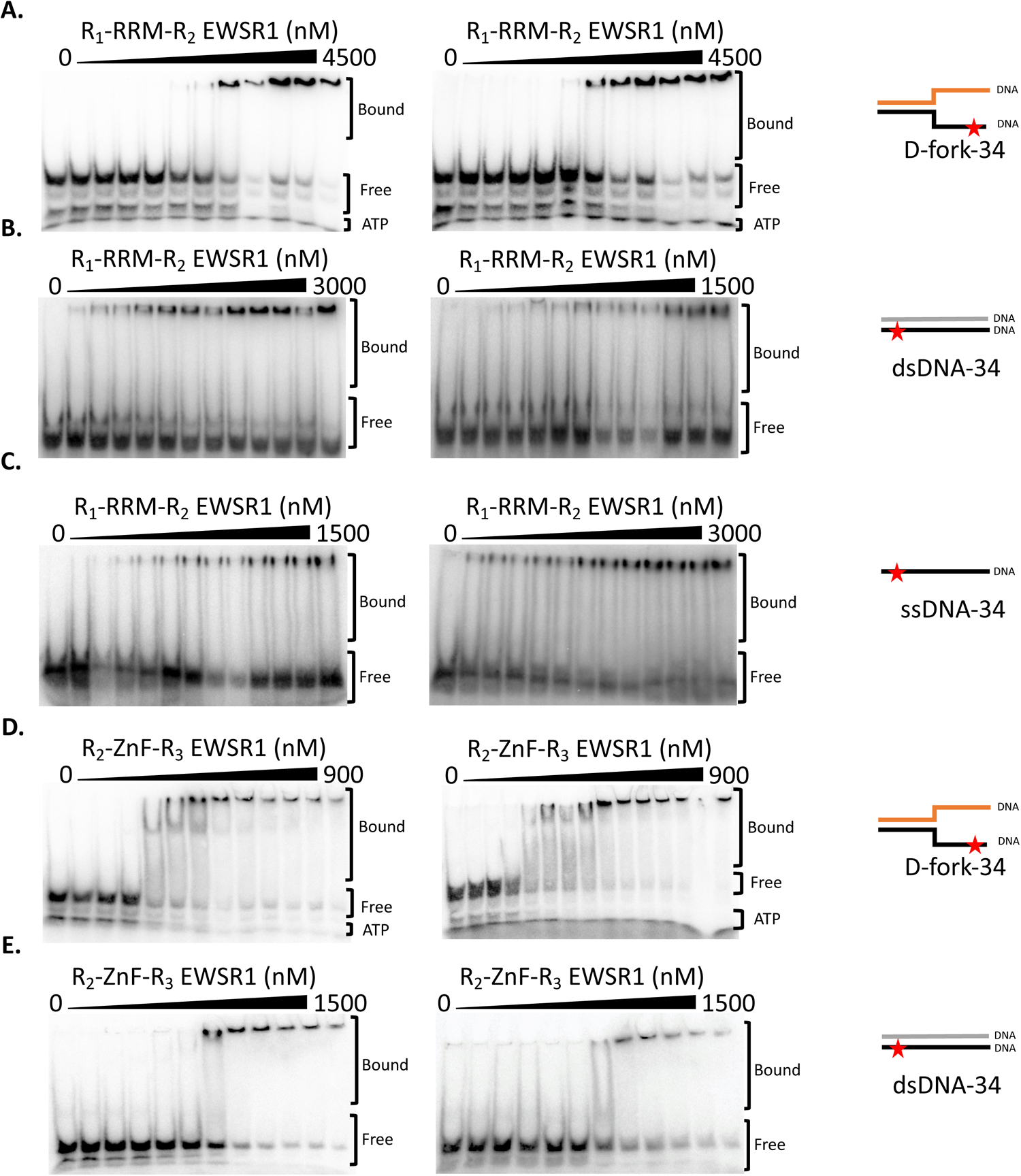
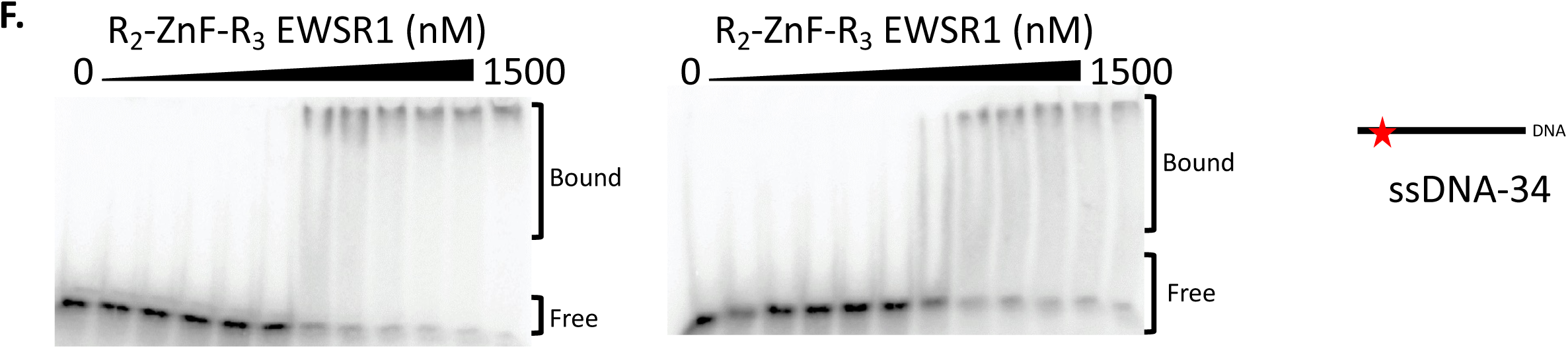
EMSA replicates for R_1_-RRM-R_2_ EWSR1 binding to (A) D-fork-34, (B) dsDNA-34, and (C) ssDNA-34, and R_2_-ZnF-R_3_ to (D) D-fork-34, (E) dsDNA-34, and (F) ssDNA-34.

